# A novel theoretical framework for simultaneous measurement of excitatory and inhibitory conductances

**DOI:** 10.1101/2021.03.30.437555

**Authors:** Daniel Müller-Komorowska, Ana Parabucki, Gal Elyasaf, Yonatan Katz, Heinz Beck, Ilan Lampl

## Abstract

Firing of neurons throughout the brain is determined by the precise relations between excitatory and inhibitory inputs and disruption of their balance underlies many psychiatric diseases. Whether or not these inputs covary over time or between repeated stimuli remains unclear due to the lack of experimental methods for measuring both inputs simultaneously. We developed a new analytical framework for instantaneous and simultaneous measurements of both the excitatory and inhibitory neuronal inputs during a single trial under current clamp recording. This can be achieved by injecting a current composed of two high frequency sinusoidal components followed by analytical extraction of the conductances. We demonstrate the ability of this method to measure both inputs in a single trial under realistic recording constraints and from morphologically realistic CA1 pyramidal model cells. Experimental implementation of our new method will facilitate the understanding of fundamental questions about the health and disease of the nervous system.

**Classification:** System Neuroscience, Cellular and Molecular Neuroscience

## Introduction

Neuronal firing is orchestrated by the interplay of excitatory and inhibitory inputs. Therefore, studying their relationship has been crucial to solving fundamental questions in cellular and system neuroscience. Disrupted relations between these inputs were suggested to accompany many neurological diseases and in particular epileptic seizures. It is commonly believed that such seizures are accompanied and even caused by a disruption of excitation-inhibition ratio and their temporal relationships ^1–3^.

The most widely used method to measure inhibitory and excitatory inputs in isolation is the voltage clamp technique. To reveal excitatory synaptic currents the membrane potential is voltage clamped near the reversal potential of inhibition (near -80 mV) and inhibitory synaptic currents are revealed when the voltage is clamped near the excitatory reversal potential (near 0 mV). Voltage clamp recordings have been used in this manner to study mechanisms of feature selectivity of cortical cells belonging to various modalities ^4–13^. Current clamp recordings also allow for the isolation of excitatory and inhibitory conductances, which is done by injecting constant positive or negative currents which bring the membrane potential near the reversal potential of these two input types ^8–10,14–18^.

Voltage and current clamp approaches share several similarities. In both cases, excitation and inhibition are recorded in different trials and conductances are estimated by fitting the averaged data with the membrane potential equation (Eq. 1 below). Hence, as these methods provide only an average picture and thus fail to capture the instantaneous and trial-by-trial based insight in the relations between excitation and inhibition.

The instantaneous relation between excitation and inhibition in-vivo were revealed using a different approach, relying on the finding that the membrane potential of nearby cortical cells in anesthetized animals is highly synchronized ^19,20^. This approach consists of depolarizing one cell in order to reveal its inhibitory inputs while simultaneously hyperpolarizing a neighboring cell to reveal its excitatory inputs. Doing this showed that excitatory and inhibitory synaptic inputs are highly correlated in anesthetized as well as in awake rodents ^21-24^ and was used to study the degree of correlation during oscillatory neuronal activities ^23^. However, this approach depends on making the recordings from highly correlated cells, mostly observed in deeply anesthetized animals. Methods for estimation of excitatory and inhibitory inputs of a single cell during single trials were developed before ^24–28^. However, these methods make important assumptions about the dynamics and statistics of the inputs. Importantly, all these methods rely on analyzing changes in membrane potential (i.e., dv/dt) when estimating excitatory and inhibitory conductances. Clearly, changes in conductance sometimes are not accompanied by any change in membrane potential, as expected when a cell receives shunting synaptic input with a reversal potential near the resting potential of the cell.

We describe a new theoretical framework for simultaneously measuring both excitatory and inhibitory conductances under current clamp in a single trial with high temporal resolution, without making statistical assumptions about the inputs. It is based on frequency analysis of the response of neurons when injected with a current composed of two sinusoidal components and allows measuring as a function of time both the excitatory and inhibitory conductances simultaneously with membrane potential. We demonstrate this method in-silico using simulations of a point neuron receiving excitatory and inhibitory synaptic inputs and then demonstrate it in a realistic pyramidal cell model when synapses are distributed further away from the soma. Finally, we describe the limitations of this approach in whole cell patch recordings obtained using contemporary intracellular amplifiers.

## Results

### Transformation of Vm a and total conductance to E and I conductances

We sought to develop a method that provides a way to simultaneously measure the excitatory and inhibitory conductances in a single trial with high temporal resolution during current clamp. We begin with the membrane equation for passive synaptic inputs (1) of a point neuron, which can be rearranged in a way that the excitatory conductance is isolated on the left (2).

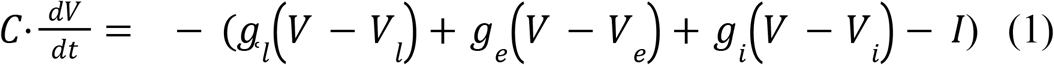

Replacing *V* − *V*_*l*_, *V* − *V*_*e*_, *V* − *V*_*i*_ with *V* ^*l*^, *V* ^*e*^, *V*^*i*^ respectively and assuming that the total conductance equals the sum of the inhibitory and excitatory conductance *g*_*s*_ = *g*_*i*_ + *g*_*e*_ we get:

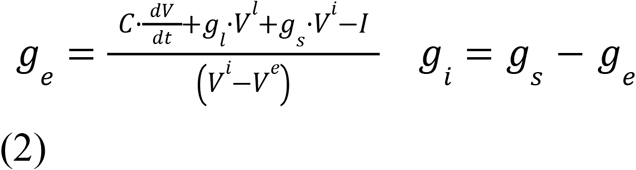

Equation (2) shows that the two inputs can be isolated if the following parameters are known: *V*, membrane voltage; *g*_*l*_, leak conductance; *g*_*s*_, total synaptic conductance; *V*_*l*_, *V*_*e*_, *V*_*i*_, equilibrium potentials of the individual conductances; *C*, membrane capacitance; *I*, stimulus current. Figure 1 shows how this equation works in a simulated point neuron where these parameters are indeed known. We demonstrate this transformation by showing depressing excitatory and inhibitory inputs as well as a step change in conductance, but it works for any type and dynamic of excitatory and inhibitory inputs.

**Fig. 1.**
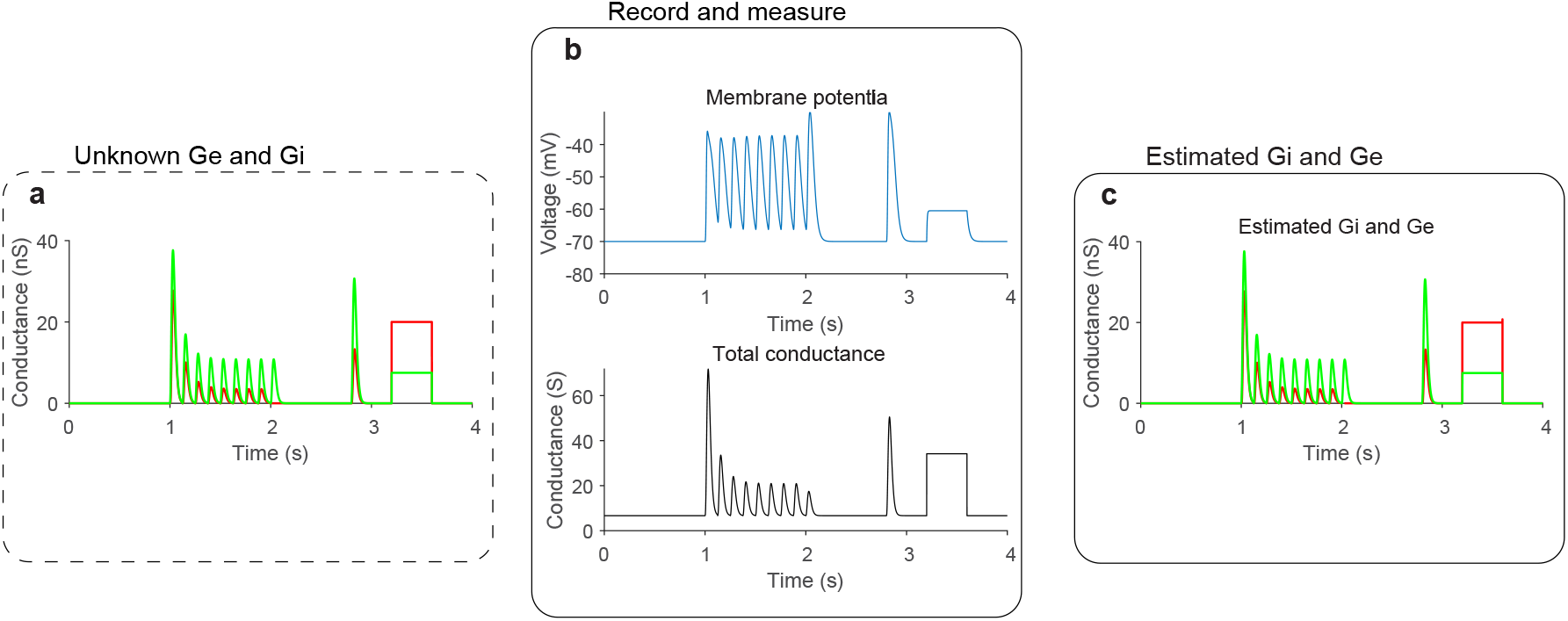
Ge and Gi can be obtained from Vm and total G. **a** Simulation of correlated excitatory and inhibitory synaptic inputs (inhibition delayed by 4 ms after excitation), which depressed according to a mathematical description of short-term synaptic depression (STD, Markram and Tsodyks, 1997). These are the inputs the method aims to reveal. **b** Membrane potential simulation of a passive point neuron (*R = 300MΩ, C =0*.*15nF*, Euler method, dt = 0.0005s) receiving the inputs in A, with the total conductance shown below. We assume that these two vectors are measurable. A short test current pulse was injected at the early part of the trace. **c** The result of transforming Vm, its derivative (not shown) and the total conductance into Ge and Gi using equation 1.

How do we find these parameters under experimental conditions? The equilibrium potentials are generally assumed to be known and determined from intracellular and extracellular ion concentrations. The leak conductance and membrane capacitance can be measured when injecting hyperpolarizing current steps. The voltage is also easy to resolve during the current clamp. However, a method to record the membrane potential and at the same time also measure the conductance at each time point has been proven to be challenging. As we describe below we can theoretically estimate the total conductance of the cell by measuring the voltage response during injection of a current composed of two high-frequency sinusoidal components. We start with impedance analysis of passive circuits representing a simplified point neuron and describing the relationships between the impedance and cell conductance.

### Impedance-Conductance relationship in a passive point neuron

To develop a method which can practically be used for whole cell patch recordings, we included the resistance of the patch pipette in our analysis. As shown below, the resistance of the electrode affects the measurement of the cell’s impedance and thus cannot be ignored. We analyzed in the frequency domain the impedance of a circuit composed of a recording electrode (Rs) and a simplified point neuron (composed of a resistor, R = 1/G, where G is the conductance of the cell, and a capacitor, C). The impedance of this circuit is given by Equation 3 (w = 2πf, i is the imaginary unit and f is the frequency in Hertz). The cell conductance (G) can vary over time, and so does the impedance of the circuit (Z).

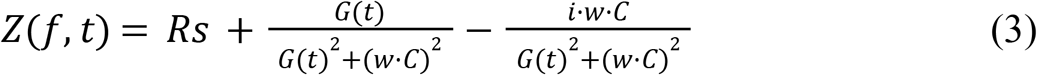

Figure 2 illustrates the relationships between the impedance and G for various frequencies. It also shows that in the presence of Rs, impedance-frequency curves intersect each other as frequency increases, resulting in a positive relationship between circuit impedance and G for a large range of G (compare Fig. 2a-c). The presence of Rs also keeps the phase almost constant for different frequencies and G values Fig. 2d). Thus the electrode resistance has a prominent effect on the total impedance of this circuit and should not be ignored when injecting high frequency sinusoidal current into cells.

**Fig. 2.**
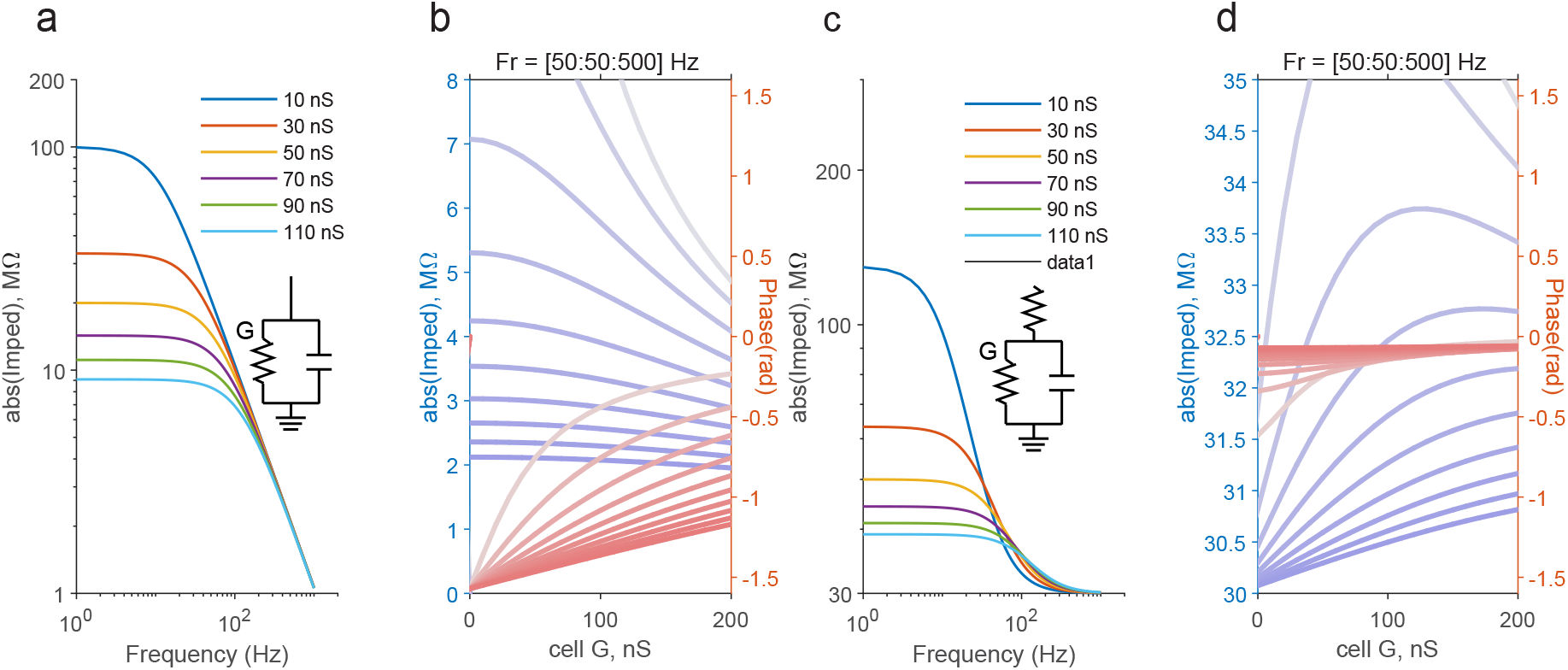
Impedance frequency-curves of passive electrical circuit for different conductances. **a** Absolute impedance as a function of frequency for different values of the model conductance. Note that none of the curves intersect., **b** Absolute impedance curves as function of conductance together with phase curves. Each line represents a different frequency (50Hz to 500Hz, steps of 50Hz, from lowest (pale blue or red) to highest (deep colors) as indicated by the text (Fr = [50:50:500]) above. **c-d** The same but when the RC circuit is also connected in series to a resistor (Rs). Note in C that curves intersect each other at high frequencies and in D that the phase is almost constant. Fixed circuit parameters: Rs = 30MΩ, C = 0.15nF.

### The in-silico experiment

In the next sections we show the response of a point neuron to an injection of a current (Fig. 3d) composed of two sinusoidal components (w1 = 2πf1, w2 = 2πf2):

**Fig. 3.**
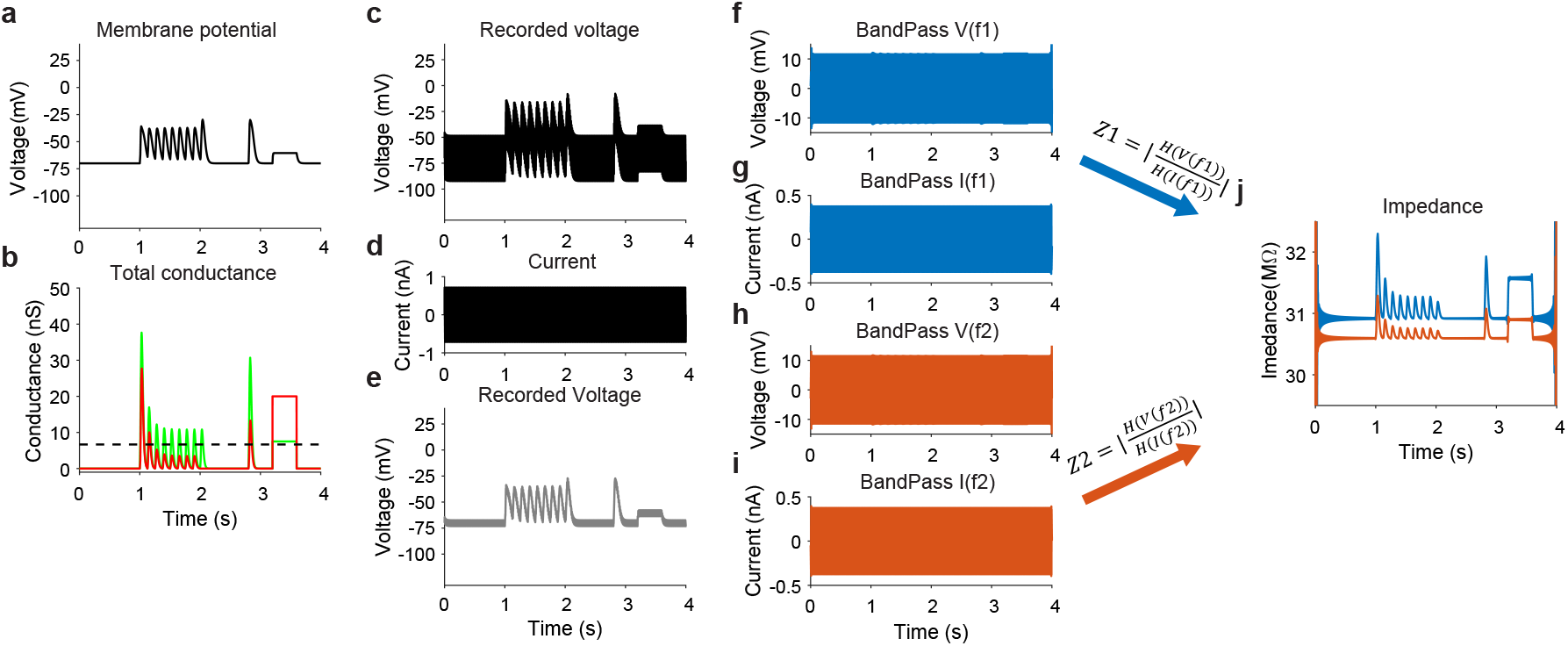
Measurement of total impedance in a single trial -simulation of a point neuron. **a**. Simulated membrane potential of a point neuron when receiving synaptic conductances as shown in **b** (excitation - green, inhibition - red). **c-d** The voltage response of a simulated neuron (c) receiving synaptic inputs described in b and injected with a current composed of two sinusoidal components (d, 0.375nA, 210Hz and 315Hz). ‘Recording’ was made via an electrode of 30MΩ and thus most of the voltage drop due to the injected current occurred across the electrode. The actual voltage change across the membrane was small (‘recorded’ when electrode resistance was set to zero). **f-i** voltage and current traces when filtered at the two frequencies used to compose the current. Note for the small fluctuations in voltage. **j** Impedance curves for each of the two frequencies obtained by dividing the Hilbert transform of the voltage and current shown in f-i and then taking the absolute values. Edge effect of the filtering is observed near zero and end times of the traces.

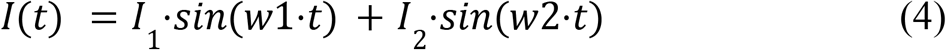

can be used to measure changes in excitatory and inhibitory conductances imposed on the model (Fig. 2b) in a single trial. Although the voltage response in our simulation fluctuates across a large range of more than 35mV (Fig. 3c), most of the drop of voltage occurs on the electrode resistor, as seen when we set Rs to zero (Fig. 3e). Across the cell itself, the fluctuations of the voltage for this example are less than 6 mV. The current and the voltage are used to calculate all the passive properties of the simulated cell in a single trial (i.e., Rs, G and C). The computations are all analytical and approximation is done only when estimating the cell’s capacitance as shown below. As described above, estimating the cell’s conductance allows to measure the excitatory and inhibitory conductances.

### Measurement of the cell total conductance

Although the experimentalis can estimate the cell’s capacitance from the response to a step current, here we show that it can be well estimated from the response to either one of the two frequencies composing the sinusoidal current (Eq. 4). We rely on the assumption that when the frequency of the current is high (w*C>>G), we can neglect G^2^ in the denominators of the second and third terms in Equation 3. Hence, at such frequencies the electrode resistance (Rs) is relatively larger than the second term, and thus the second term can be neglected. In this case, the total impedance of the circuit is mostly determined by the electrode resistance and the capacitance of the cell, as the latter draws most of the sinusoidal current that is injected into the cell. Hence, the phase relationship between the voltage and the current can be approximated by equation 5 (see also the phase curves in Fig. 2d).

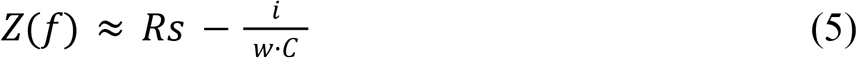

For such estimation to be valid, the frequency of each one of the two current components has to be sufficiently high. For example, for a cell with a mean conductance of 1/100MΩ and total capacitance of 0.15nF, recorded with 10MΩ electrode (Rs), a ratio of ∼88 between (w*C)^2^ and G^2^ will be obtained at 100Hz. Since the impedance of the second term in Equation 3 for this example is ∼1MΩ, much smaller than Rs (10MΩ), we neglect this term. Thus, the capacitance can be obtained from Equation 5, if we can estimate the electrode resistance and the phase relationship between the current and the voltage. We do this in a single trial by first measuring the electrode resistance (Rs_est_) from the ratio of the absolute values of the fast Fourier Transform (FFT) of the voltage and the current at the frequency of the injected current, after both traces were bandpass filtered at one of the two frequencies (f1 or f2, using ‘bandpass’ Matlab function, implementing finite impulse response (**FIR**) filter). The two vectors (FV,FI bandpass filtered voltage and current) are then used to estimate Rs:

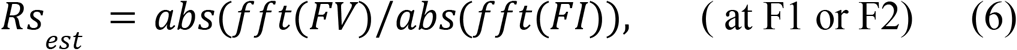

The phase between FV and FI is calculated from the Hilbert transform of FV (H operator, either for the F1 or F2) using the ‘hilbert’ Matlab function and averaging over time:

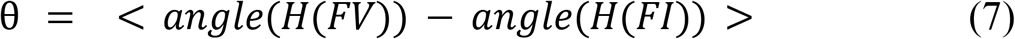

Equations 5-7 allow estimation of the cell’s total capacitance:

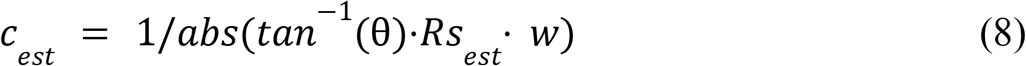

In the example shown in Figures 2 and 3, the real capacitance was set to 0.15nF and was estimated as 0.149nF. Note, that estimation of C can also be obtained when setting Rs to zero at a similar accuracy.

We then use the estimated capacitance of the cell to to measure the cell’s conductance and to obtain a more accurate measurement of the electrode resistance, both over time in a single trial. In this computation these values will be measured based on the analytical solution of Equation 3, this time without making any approximations. Here we use the fact that the current contains two sinusoidal components having two different frequencies (F1 and F2, e.g., 210Hz and 315Hz as used in the example). Since Z(f) decreases with increasing frequency (Fig. 2), increasing the frequencies, although allows higher temporal resolution, will reduce the signal to noise ratio in the presence of noise. The voltage and the current are then bandpassed at the two frequencies (Figs. 3f to i, due to screen resolution are as patches of colors). Note for the small modulations in the bandpassed voltage signals, which are in the order of about 1mV. These modulations result from changes in the cell’s conductance during the simulation of the synaptic inputs following the relationships between them as shown in Fig. 2). For each voltage and current bandpassed traces, ‘FV1(t)’, ‘FV2(t)’, ‘FI1(t)’, ‘FI2(t)’, we then compute the hilbert transforms (HeVF1(t), HeVF2(t), HeIF1(t), HeIF2(t), using the ‘hilbert’ Matlab function). These complex vectors are then used to calculate the impedance of the cell at the two frequencies:

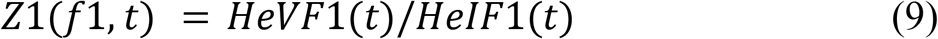

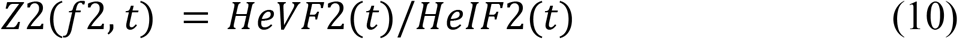

The absolute values of these complex vectors, shown in Figure 3j, demonstrate curves with a shape that is similar to that of the total conductance of the cell (leak plus synaptic conductances). Note that when the coductantance of the cell is increased during activation of these inputs, the impedance is also elevated. This only happens in the presence of Rs, as shown in Fig. 2.

These two impedance vectors are then used together to solve Equation 3 and obtaining a solution for Rs and G (when *z*1 ≠ *z*2, c is the estimated capacitance). To this end we used Mathematica (Wolfram) to solve the two equations for absolute values of z1 and z2 (“**Solve[Abs (r + 1/(g + I*w1*c)) == Abs (z1) && Abs (r + 1/(g + I*w2*c)) == Abs (z2), {r, g}]”, I = imaginary unit)** which gives the following solutions for Rs and G (here Z1 and Z2 are complex time dependent vectors):

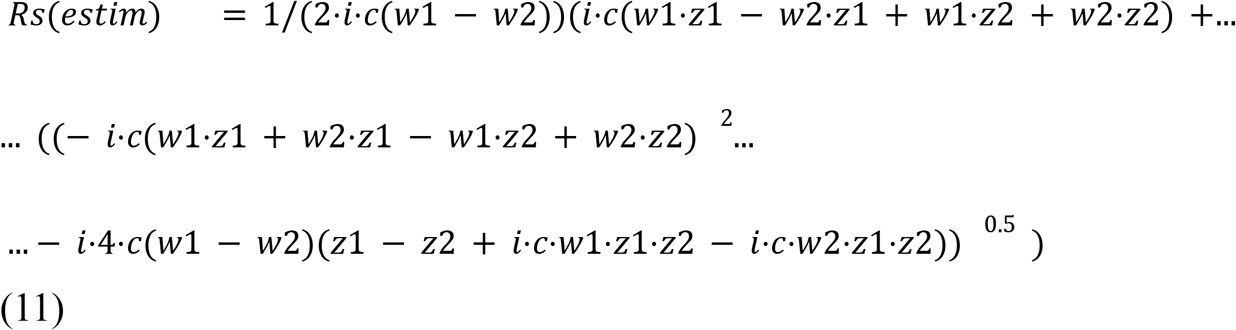

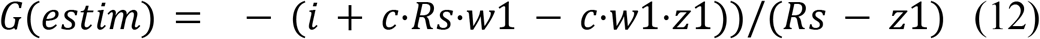

In equations 11 and 12 *Rs*_*est2*_and *G*_*est2*_ are time dependent variables. In Figure 4a we again plotted the two impedance curves and included also the electrode resistance (Rs), which is only slightly larger than its real value used in the simulation. The estimated total conductance is plotted in Figure 4c. Note that the estimated total conductance is almost identical both in shape and magnitude to the sum of the leak, excitatory and inhibitory conductances used to simulate the membrane potential in this example (*G*(*estim*) = *g*_*l*(*estim*)_ +*g* _*s*(*estim*)_).

**Fig. 4.**
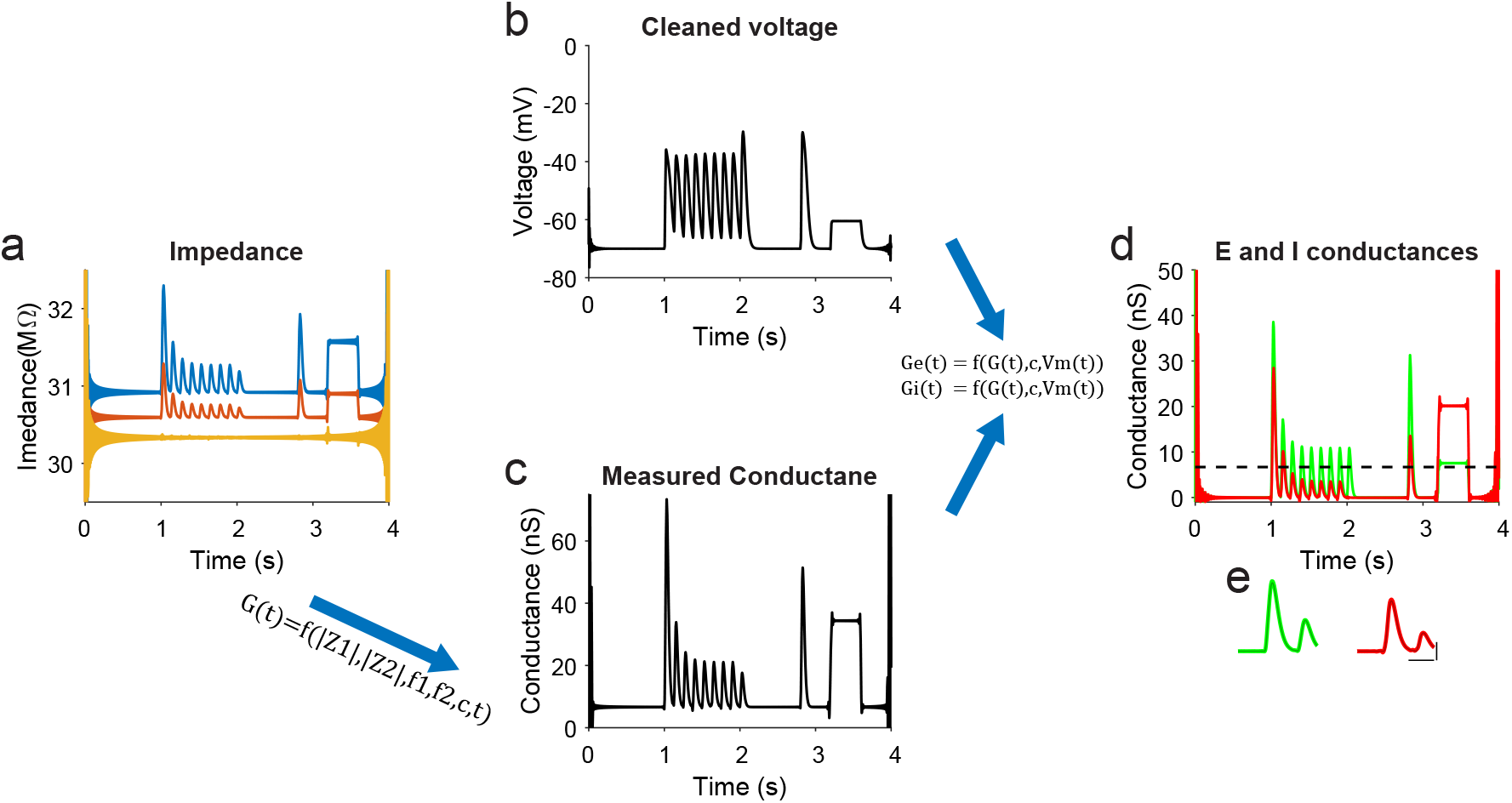
Measurement of Ge and Gi in a single trial from impedance measurements - simulation of a point neuron. **a** Absolute impedance curves for each frequency (as shown in Fig. 2). **b** The membrane potential after filtering out the components at the injected frequencies. **c**, Measured conductance and electrode resistance (shown in A) were obtained from the absolute impedance curves (Eq. 12, 13) and the estimated capacitance. **d** Excitatory and inhibitory conductances were estimated from the cleaned Vm and measured conductance (Eq. 1,2). **e** Extended data for the rarely synaptic responses, superimposed with the real conductances (thin lines) shown in D (scale bars are 100 ms and 10nS.

### Estimation of the excitatory and inhibitory conductances from cell’s conductance and membrane potential

After estimating the total conductance, Equations 1 and 2 are used to compute E and I conductances as discussed above. Since sinusoidal current is injected into the cell (with two frequency components) we remove these signals from the voltage trace to obtain a clean version of the membrane potential. Before we use equations 1 and 2, we need to calculate the resting membrane potential and its corresponding leak conductance. We do it by finding the mean voltage in the cleaned membrane potential for the lower 5^th^ percentile of the total conductance vector, which we assume reflects the resting state at which no synaptic inputs are evoked (i.e, gl). The corresponding membrane potential values for this 5th percentile conductance were used to calculate the mean resting potential (i.e.,V_1_). The synaptic conductance is simply given by *g* _*s*(*estim*)_ = *G*(*estim*)−*g* _*l*(*estim*)_. In the transformation presented in equation 1 and 2 we assume that the reversal potentials of excitation and inhibition are available to us. The capacitance and total conductance are obtained as described above. The results of these computations are shown in Figure 4d. Our calculations revealed that the estimated conductances are almost identical to the real inputs of the simulated cell (compare Fig. 3b to 4d). We note that our method allows estimating the conductances even when tonic input exits, as demonstrated in the step change in E and I (shown between 3 to 4 seconds). In fact, the Pearson correlation between the real inputs and the estimated inputs for this simulated example were extremely high: 0.999 for excitation and 0.996 for inhibition (not shown).

### Computing the excitatory and inhibitory conductances of a cell embedded in a balanced network

We asked if our approach can be used to reveal the underlying excitatory and inhibitory conductances of a model cortical neuron embedded in an active network where it receives excitatory and inhibitory inputs. Therefore we used a simulation of a cortical network at a balanced asynchronous state ^29^ to obtain the excitatory and inhibitory synaptic inputs of a single cell (kindly provided by Dr. Michael Okun, University of Leicester). We used these conductances in a simulation of a single cell, in which we injected a current with two sinusoidal components (210 Hz and 315Hz) via a 50MΩ electrode and measure the response of the cell, before (Fig. 5) and after filtering out the two sinusoidal components from the membrane potential (Fig. 5b, black trace, which is superimposed almost perfectly with the one obtained without current injection, blue trace).

**Fig. 5.**
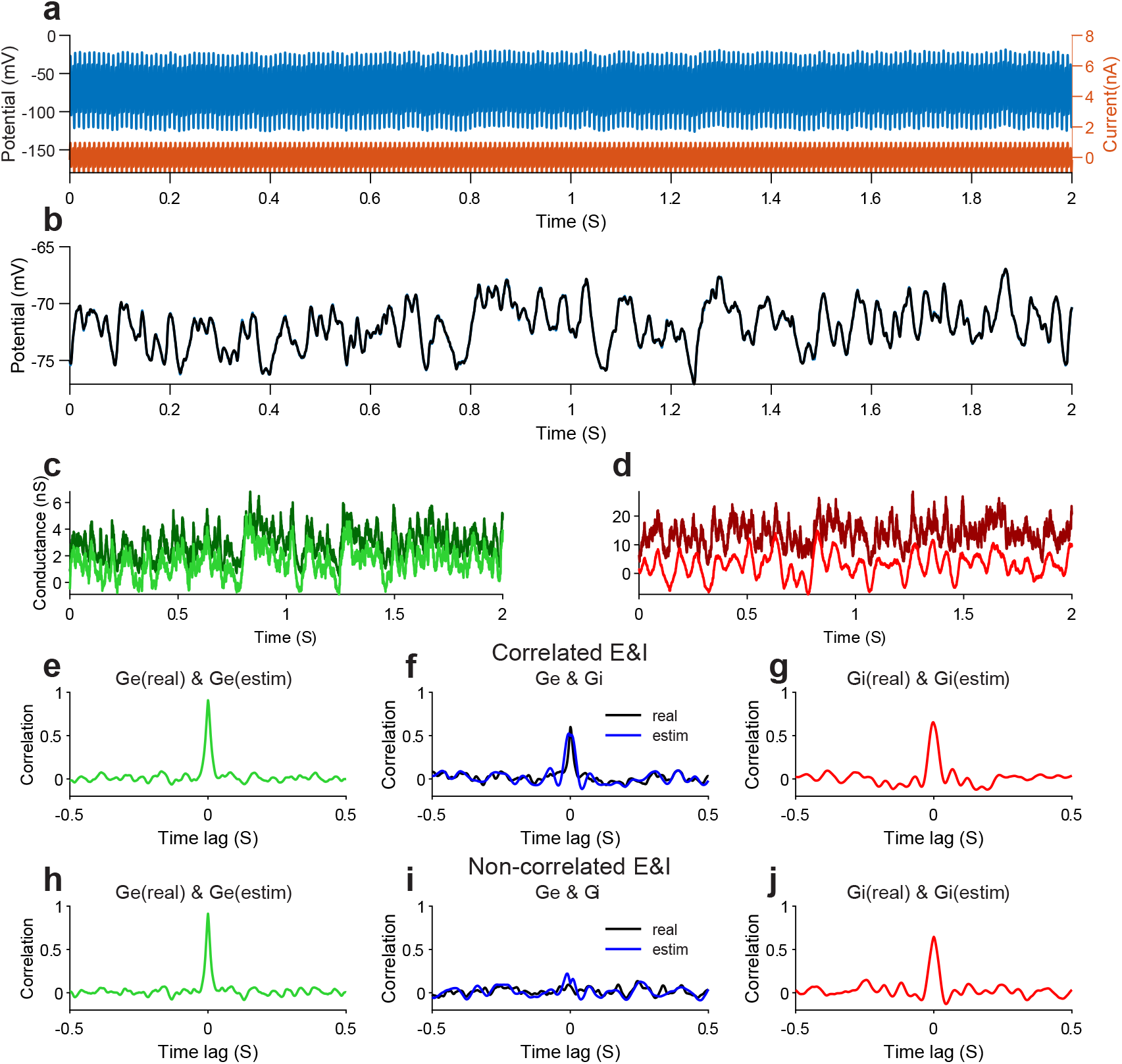
Estimation of excitatory and inhibitory conductances of a neuron embedded in balanced/unbalanced networks. **a** Excitatory and inhibitory conductances were used to simulate the response of a point neuron recorded with a 50MΩ pipette while injected with 210Hz & 315Hz 0.5 nA sinosiodal current. **b** The voltage in A was ‘cleaned’ from the sinusoidal component (black trace) and it is displayed with the voltage response of the cell when no current was injected superimposed with the ‘cleaned’ voltage. **c-d** Estimated excitatory and inhibitory conductances superimposed with the imposed conductances of the simulated cell. **e-f** Color lines describe the cross-correlations between the imposed and estimated conductances and the black lines are the correlations between imposed (real) Ge and Gi, which are lower than the real-measured correlations for each input.

We then used our computations to estimate the excitatory and inhibitory conductances (Fig. 5c and D). Note, however, that for both inputs the estimated conductances are more negative than expected. This is simply because the leak conductance was estimated from the 5^th^ percentile of the total conductance of the cell, but since synaptic activity persisted throughout the trace, the leak conductance reflects a mixture of the true leak conductance and some baseline synaptic activity. Nevertheless, the estimated excitatory and inhibitory synaptic conductances were very similar to those used as inputs (Fig. 5e,g) and similarly to the real inputs, estimated E and I conductances were highly correlated (Fig. 5f). Our approach was also successful in measuring E and I inputs when they are not correlated (Fig. 5h, j, by shifting the inhibitory input by 10 seconds relative to excitation). Indeed, as expected for this case, no correlation was measured between the measured inputs (Fig. 5i). In summary, our approach allows accurate estimation of excitatory and inhibitory inputs in various conditions without any need to take into account the dynamic and statistical properties of the excitatory and inhibitory inputs.

### Measurement of E and I inputs during large variations in access resistance

Changes in access resistance due to incompletely ruptured membrane or other due to movement of the recorded cell and preparation, pressing the pipette onto the membrane, are well known limitations of whole cell patch recordings. However, one of the advantages of the approach is in its ability to track changes in the electrode and access resistance and taking them into account when calculating the total conductance of the cell with a high temporal resolution. We demonstrate it by simulating rapid changes in the electrode resistance during the in-silico recordings (Fig.6, identical synaptic inputs to those used in Fig. 3). These variations led to noisy impedance measurement (Fig. 6a). However since we can measure Rs and total G at the same time (Eqs. 11-12), the changes in the electrode resistance had no effect on the ability to accurately estimate the inhibitory and excitatory conductances (Fig. 6d).

**Fig. 6.**
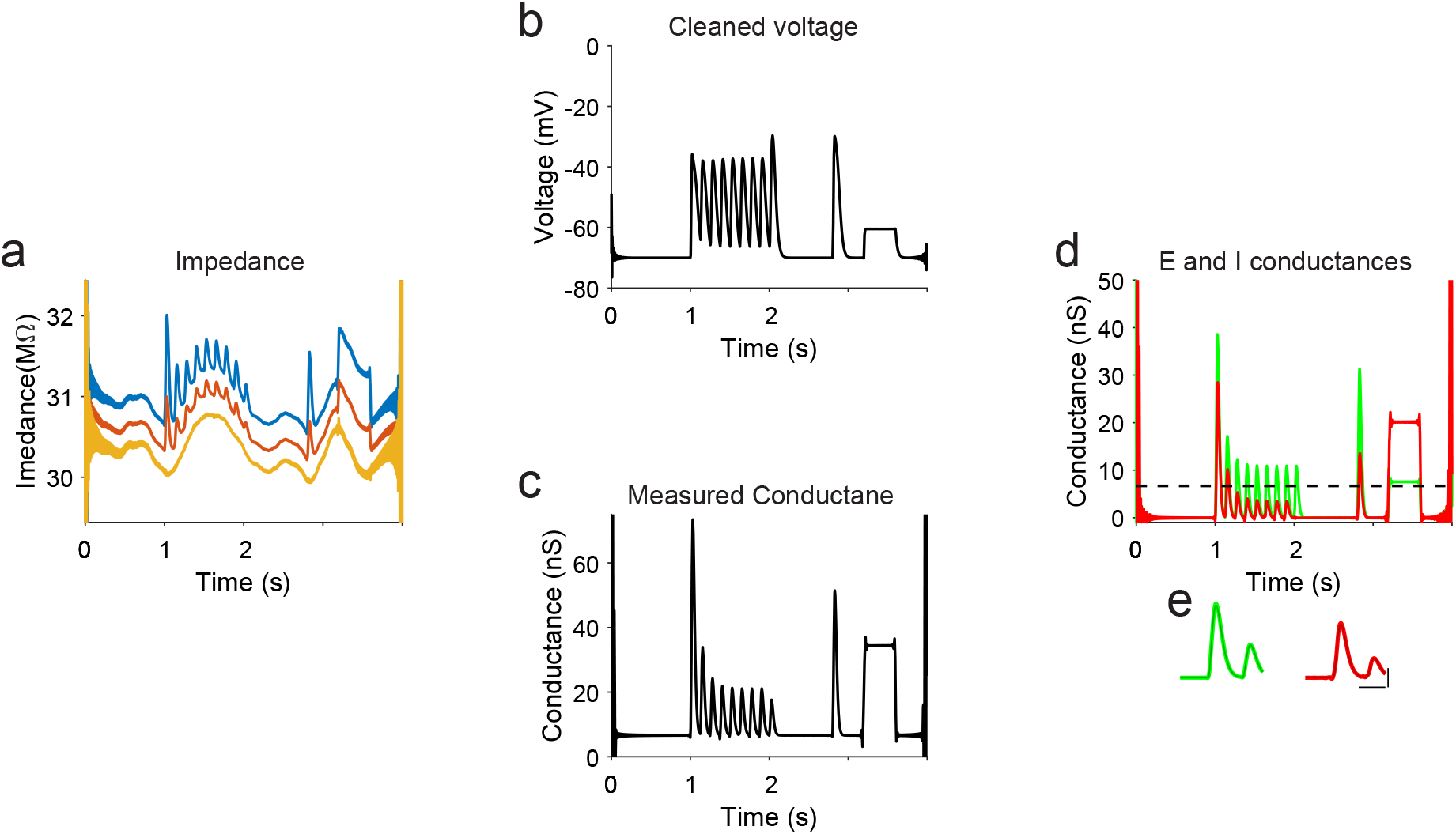
Measurements of Ge and Gi are accurate even when electrode resistance is not stable. **a** Absolute impedance curves for each frequency (as shown in Figure 2) depicted together with measured electrode resistance. Electrode resistance was modulated by smoothing lowpassong Normally distributed noise. All other parameters of the inputs are identical to those shown in Figures 3-4. **b-d**. Analysis of the voltage response of the point neuron as processed by the same way as shown in Figures 2-3. Note for the accurate estimates of E and I conductances (compare to Fig. 3b).

### Compensation for electrode capacitance

In the above computations we assumed that the recordings are made with a pipette of zero capacitance. However, electrode capacitance can greatly affect the measurement using our novel algorithm. Most of the stray capacitance of recording pipettes is formed by the separation of the solution inside the glass pipettes and that outside the pipette. Experimentally it can be reduced but not eliminated by coating the pipette with hydrophobic material ^30^. Pipette capacitance (Cp, illustrated in Fig. 7) can be also neutralized by the electronic circuit of the intracellular amplifier, using a positive feedback circuit. In our in-silico experiment we show that Cp can greatly affect the measurement, as pipette capacitance draws some of the injected sinusoidal current. As a result, the impedance measurements for the two frequencies (z1 and z2) are smaller than expected from the cell and Rs alone (Fig. 7b, Rs is 20MΩ and the curves are well below this value). This in turn results with much higher leak conductance and a completely wrong estimation in the synaptic conductances based on equations 11-12. All together, our estimations can be flawed, leading to negative evoked inhibitory conductance (Figure 7d).

**Fig. 7.**
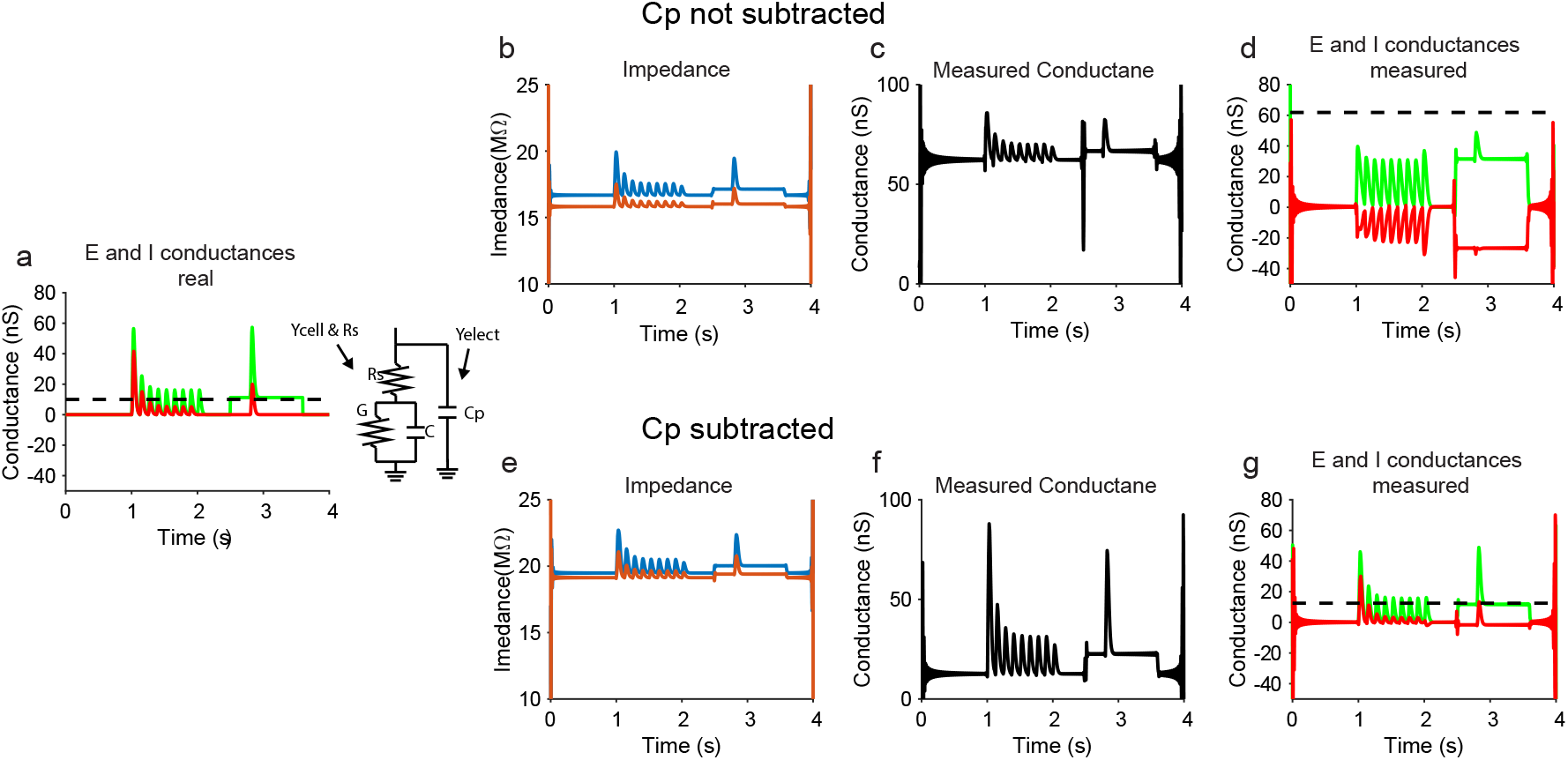
Computational approach for compensation of parasitic capacitance for measurement of excitation and inhibition in an in-silico model. **a** The ‘real’ inputs in the in-silico experiment and the circuit representing the recording configuration. Note for Cp, the parasitic capacitance of the electrode (Rs). **b-d** Measurement of the impedance, followed by calculation of the conductance and E and I inputs using the approach described in Figures 3-4. **e-g** Similar measurements when recalculating the impedance at each frequency using equations 14-18 to subtract the effect of pipette capacitance.

To compensate for the reduction of the impedance due to the pipette capacitance we estimated Cp and then used this value to correct the measured impedances. Here we calculate the admittance (Y, Y = 1/Z) at each of the two frequencies for the equivalent circuit of a cell recorded with a pipette that has stray capacitance, as shown in Figure 7. The second terms in the following equations depict the admittance of the stray capacitance.

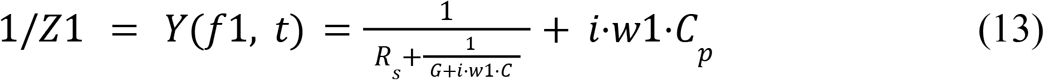

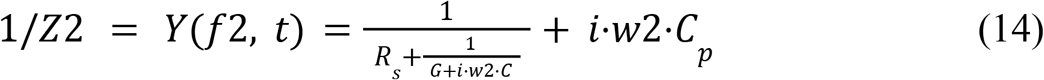

From these two equations and replacing 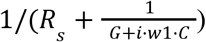 with Y1 (and Y2), C_p_ is given by:

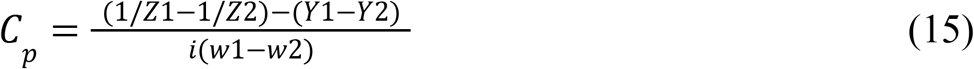

However the value of Y1 and Y2 are unknown and are those we seek. We found, however, that the second term (Y1-Y2) can be neglected as it is much smaller when compared to the value of (1/*Z*1−1/*Z*2). For example, for the parameters used in this simulation, the ratio between the latter and first terms is ∼200, clearly justifying our next approximation:

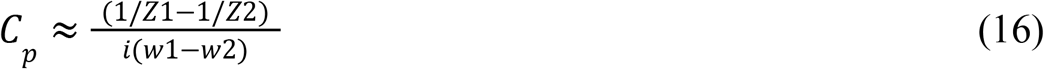

We then use this estimated value of C_P_ to calculate the estimated impedance of the cell and the electrode alone 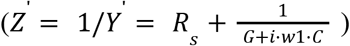 which is one by subtracting from the measure Z’ the C_P_ component following rearranging equations 13 and 14:

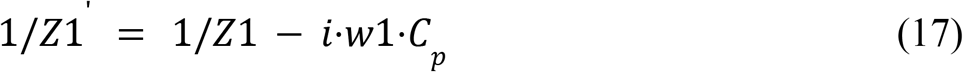

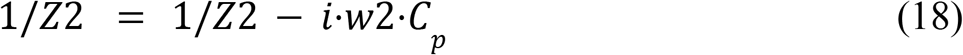

The new Z’ vectors are then used as the inputs as described above in equations (11-12) and the subsequent process. This approach greatly improved the measurement of excitation and inhibition (Figs. 7e-g). Hence, this component in the analysis, which can be switched on and off, can help in resolving the analysis of real recordings, where stray capacitance always exists.

### Measuring Dendritic Conductances

To assess how our method resolves dendritic conductances, we simulated a morphologically realistic CA1 pyramidal cell ^31^. We uniformly distribute 50 inhibitory and 50 excitatory synapses proximal to the soma (Fig. 8a). We realized that due to current escape of the injected sinusoidal current to the dendrites, the estimated leak conductance is much larger than its actual value. In the case of proximal synaptic inputs, less current is escaping towards the dendrites during activation of these inputs when compared to pre-stimulation conditions. We compensated for this change by dynamically altering the strength of the leak conductance at each time point based on the estimated total synaptic conductance before calculating the excitatory and inhibitory conductances (Eq. 1-2) by using this empirical equation:

**Fig. 8.**
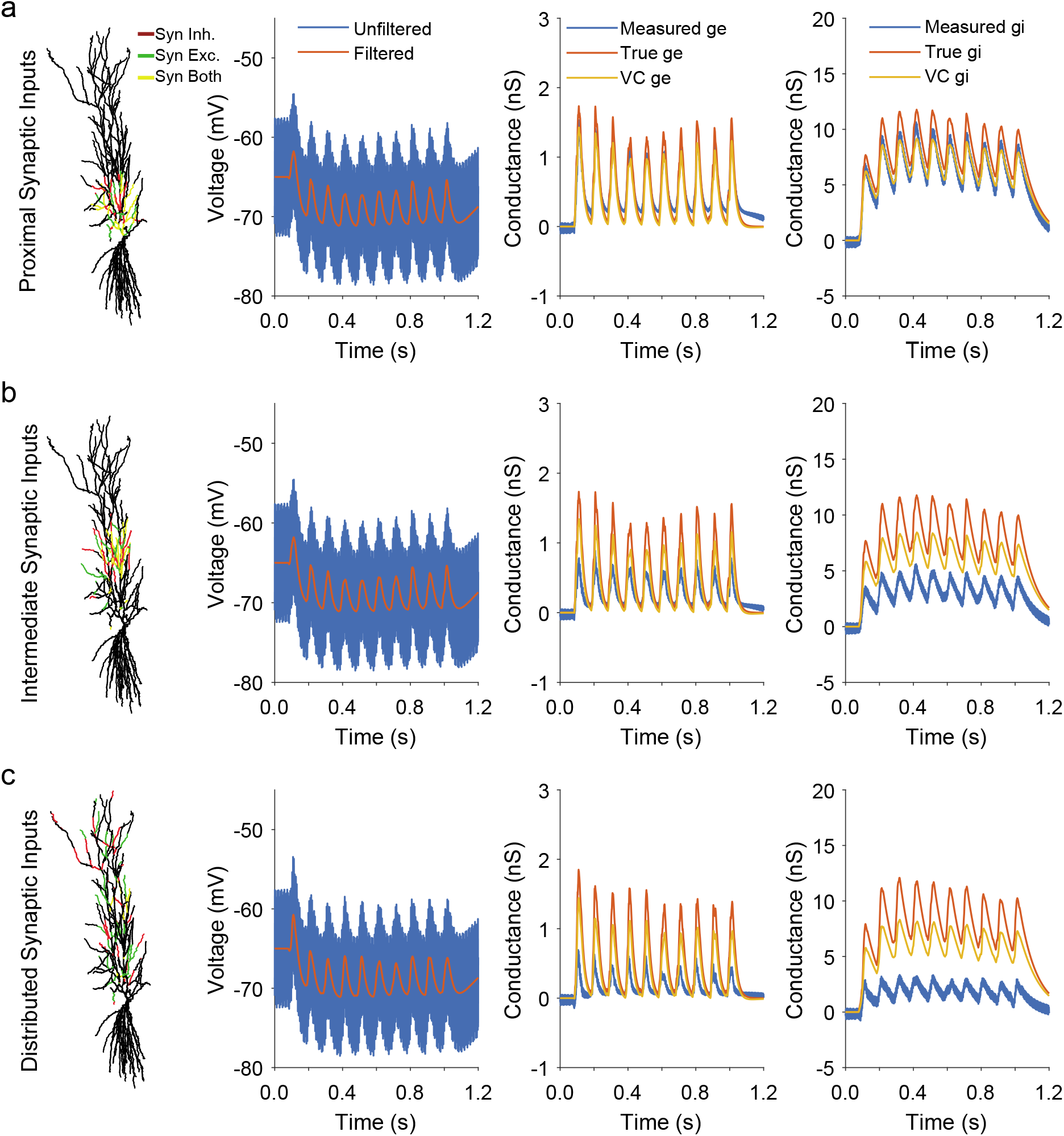
Measuring conductance changes of dendritic synapses. **a** Inhibitory and excitatory synapses were placed proximal (129.92µm ±47.83µm STD) to the soma and all measurements were performed at the soma. Simultaneous conductance measurements are at least as accurate as separate voltage clamp recordings for these proximal synapses. **b** Inhibitory and excitatory synapses were placed at intermediate distance (238.69µm ±39.71µm STD). Simultaneous conductance measurements and voltage clamp are both well correlated with the temporal dynamics but underestimate the amplitude. Simultaneous conductance measurements underestimate amplitude to a greater extent. **c** For distributed synapses (309.92µm ±164.46µm STD) simultaneous conductance measurements become less accurate but still follow the true conductances.

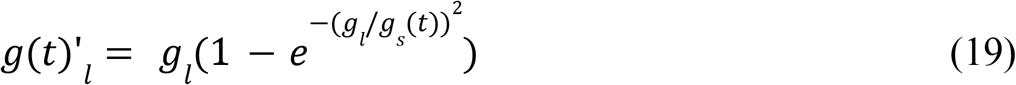

It shows that for weak proximal synaptic input this function strongly reduces the newly calculated leak conductance (g(t)’) as expected, and that this allows to compensate for the current escape. Another way to look at it is as if the electrotonic length of the cell was small at this condition. However, when the synaptic inputs get stronger the function increases the leak, as less current is expected already to escape to the dendrites due to the shunting effect of the input.

Although those synapses are on average 129.92µm (±47.83µm STD) away from the soma, our method resolves the excitatory and inhibitory conductances in a single trial, at least as well as the voltage clamp measurements done during two separate trials. When the synapses are moved further away, to an intermediate distance of 238.69µm (±39.71µm STD), our method underestimates the conductances to a larger extent than voltage clamp (Figure 8b). Under most biological conditions synapses are not constrained to a narrow part of the dendrite. Therefore we uniformly distributed synapses anywhere on the apical dendritic tree (Fig. 8c). This resulted in synapses with an average distance to the soma of 309.92µm (±164.46µm STD). In this case our method still follows the conductances but underperforms as compared to voltage clamp. Because the measurement quality seemed to decrease with distance, we did more simulations to quantify the relationship between somatic distance and recording quality.

### Proximal Conductance Measurements are Stable and Reliable

To investigate the relationship between measurement quality and synaptic distance to soma, we simulated a single excitatory and a single inhibitory synapse at the same dendritic segment. As above, we found that we can reliably isolate the conductances when the synapse pair is close to the soma (Figure 9a). At an extremely distal point the measurement becomes unreliable. Even the voltage clamp ceases to follow the temporal dynamics but it is at least positively correlated with the true conductance. To quantify the extent to which our measurement follows the temporal dynamics of the current we calculated the correlation coefficient between measurement and true conductance. We found that the measurements are extremely reliable for synapses below 300µm somatic distance (Figure 9c). Above that distance, the measurement quality breaks down abruptly, eventually becoming negatively correlated (Figure 9b,c).

**Fig. 9.**
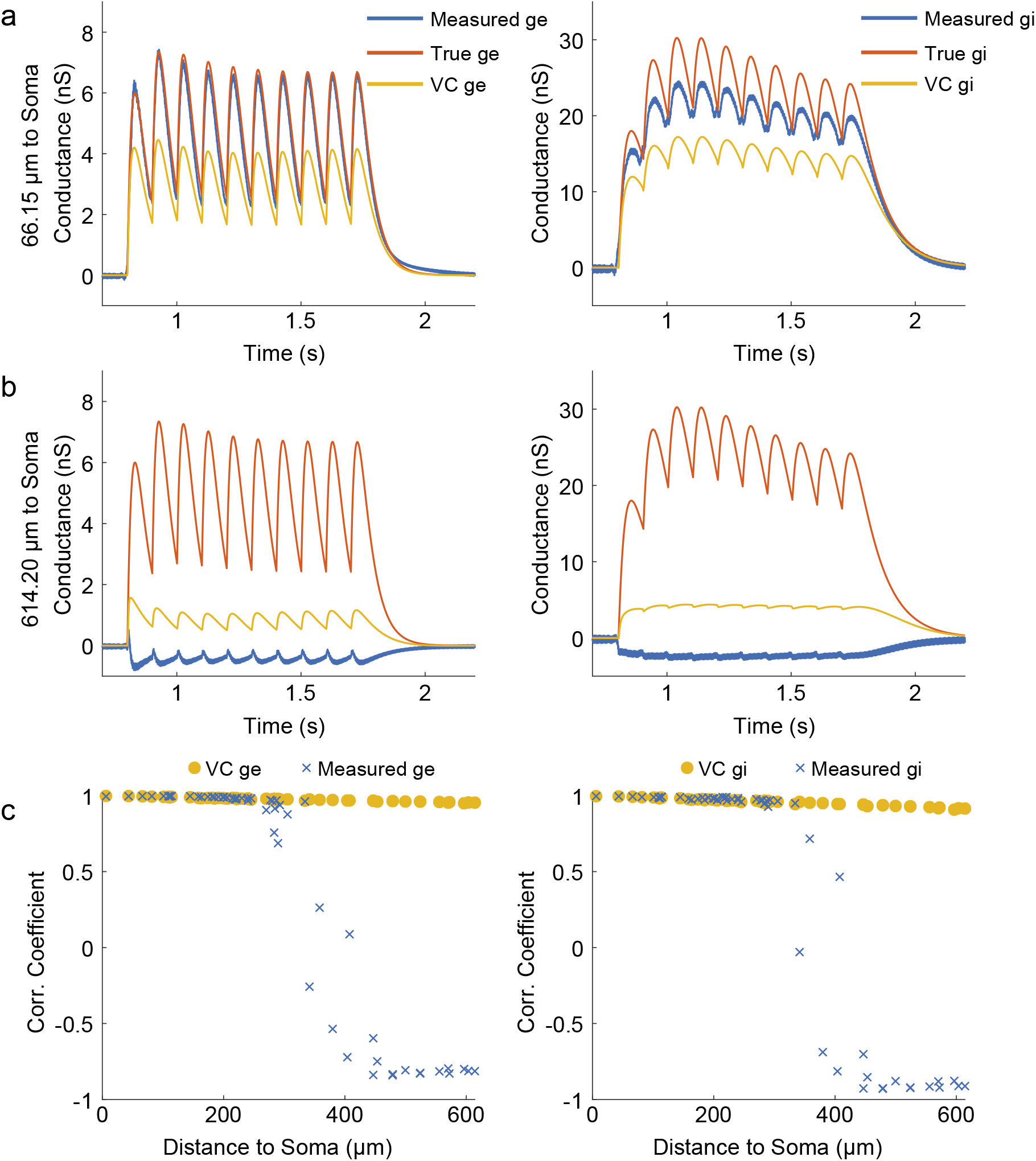
Simultaneous conductance measurements are accurate for synapses up to 300µm but break down above. **a** Simultaneous conductance measurements are highly accurate for the excitatory and inhibitory proximal synapses. **b** For extremely distal synapses our simultaneous conductance measurements become inaccurate. Color legend as above. **c**,The correlation coefficient for many synapses confirms that simultaneous conductance measurements are at least as accurate as voltage clamp below 300*µm somatic distance. For further away synapses the conductance measurement technique breaks down abruptly*.

## Discussion

We describe a novel framework to estimate the excitatory and inhibitory synaptic conductances of a neuron under current clamp in a single trial with high temporal resolution while tracking the trajectory of the membrane potential. We show that the method allows estimating these inputs also in a morphologically realistic model of a neuron. The work described above here is theoretical and lays the foundations for future experimental work.

The method is based on the theory of electrical circuit analysis over time when a cell is injected with a sum of two sinusoidal currents. This allows to measure excitatory and inhibitory conductances and at the same time track the membrane potential.

We demonstrated the method in simulations of a point neuron and in realistic simulations of a pyramidal cell, receiving proximal and uniformly distributed synaptic inputs. For the point neuron, we showed that we can reveal the timing and magnitude of depressing excitatory and inhibitory synaptic inputs with high temporal resolution and accuracy of above 99% (Figs. 3 and 4). In another example, we used our method to reveal these inputs during an asynchronous balanced cortical state and show that excitation and inhibition dynamics can be measured with high accuracy. Importantly, these estimations were obtained at a single trial and allowed obtaining the natural dynamics of the membrane potential by filtering out the sinusoidal components of the response to the injected current. Therefore, our method is suitable for estimation of excitation and inhibition in particular when these inputs are not locked to stereotypical external or internal events, such as during ongoing activity. We note that when injecting high frequency current (of a couple of hundred Hertz and above), the voltage drops mostly across the recording electrode. Here we tuned the current amplitude to produce a few millivolts sinusoidal fluctuation across the cell membrane, which should have minimal effect on voltage dependent intrinsic and synaptic conductances when performing recordings in real neurons.

### Comparisons with other methods

#### Measurement of average excitatory and inhibitory conductances of single cells

Excitatory and inhibitory synaptic conductances of a single cell were measured both under voltage clamp or current clamp recordings, focusing *in-vivo* on the underlying mechanisms of feature selectivity in sensory response of cortical cells and on the role of inhibition in shaping the tuned sensory response of mammalian cortical neurons ^6,8,35^. Conductance measurement methods were also used to reveal the underlying excitatory and inhibitory conductances during ongoing Up and Down membrane potential fluctuations, which characterize slow wave sleep activity ^36,37^. The advantages and caveats of these methods were reviewed in ^38^. Common to these conductance measurement methods is the requirement to average the data over multiple repeats, triggered on a stereotypical event (such as the time of sensory stimulation or the rising phase of an Up state) and then average trials at different holding potentials. The averaged data is then fitted with the membrane potential equation (assuming that the reversal potentials are known) to reveal the conductance of excitation and inhibition at each time point. However, these methods cannot reveal inhibition and excitation simultaneously in a single trial and only estimate averaged relationships. Our proposed method, on the other hand, allows for simultaneous measurements at a single trial. Importantly, since there is no need to depolarize or hyperpolarize the cell, our method allows measurement of synaptic conductances at the resting potential of the cell, potentially obtaining measurements of voltage dependent conductances as they progress during the voltage response to the synaptic inputs.

An alternative approach for estimating the excitatory and inhibitory conductances of a single cell was demonstrated for retinal ganglion cells ^39^. In this study the clamped voltage was alternated between the reversal potential of excitation and inhibition at a rate of 50 Hz and the current was measured at the end of each step. This study revealed strong correlated noise in the strength of both types of synaptic inputs. However, unlike the method proposed here, the underlying conductances are not revealed simultaneously and due to the clamping, the natural dynamics of the membrane potential is completely unavailable, preventing examining the role of intrinsic voltage dependent dynamics in the generation of neuronal subthreshold activity.

#### Single trial measurements of E and I under various assumptions on synaptic dynamics

Theoretical and experimental approaches based on the dynamic of excitatory and inhibitory conductances in a single trial were proposed before. Accordingly, excitation and inhibition are revealed from current clamp recordings in which no current is injected. Approaches based on Baysian methods which exploit multiple recorded trials were proposed ^25^ and estimation of these inputs in a single trial were also proposed but lack the ability to track fast changes in these conductances ^24^. A group of other computational methods ^26–28^ showed that excitatory and inhibitory conductances can be revealed in a single trial when analysing the membrane potential and its distribution. Common to all these methods, is the requirement to observe clear fluctuations in the membrane potential. Our method, however, allows revealing these inputs even if no change in membrane potential due to synaptic input is observed (except for the response to the injected sinusoidal current). Changes in conductance are often expected even when the membrane potential is stable, for example when a cell is receiving tonic input (see the step change in excitation and inhibition in Figs. 3 and 4 (between 3 to 4 seconds) resulting in a constant membrane potential value) and when a constant balance in excitatory and inhibitory currents exists.

#### Paired intracellular recordings

The substantial synchrony of the synaptic inputs among nearby cortical cells ^19,21,40,41^ allows continuously monitoring both the excitatory and inhibitory activities in the local network during ongoing and evoked activities. A similar approach was also used to study the relationships between these inputs in the visual cortex of awake mice ^22^ as well as gamma activity in slices ^4^. While paired recordings are powerful when examining the relationships between these inputs in the local network, such recordings do not provide definitive information about the inputs of a single cell. Moreover, although the instantaneous relationship between excitatory and inhibitory inputs can be revealed by this paired recording approach, the maximum inferred degree of estimated correlation between excitation and inhibition is bounded by the amount of correlation between the cells for each input, which may change across stimulation conditions or brain-state ^42–44^. For example, a reduction in the correlation between excitation, as measured in one cell, and inhibition measured in the other cell can truly suggest smaller correlation between these inputs for each cell, but it can also result from a reduction of synchrony between cells, without any change in the degree of correlation between excitation and inhibition of each cell. This caveat of paired recordings prevents us from finding, for example, if cortical activity shifts between balanced and unbalanced states ^45,46^. Simultaneous measurement of excitatory and inhibitory conductances of a single cell across states will allow these and other questions to be addressed.

### Limitations

Theoretically, increasing the frequency of the sinusoidal waveforms of the injected current in our method improves the temporal precision when measuring synaptic conductances. However, this comes at the expense of sensitivity, which reduces as frequency increases (Eq. 3 and Fig. 2). In our simulations we limited the frequency of the injected current up to about 350Hz. At this range, our simulations, depicting realistic passive cellular properties and typical sensory evoked conductance will result with a clear modulation in voltage when injecting ∼1nA sinusoidal current. When bandpass filtering the voltage, the modulation is small, in the order of a millivolt, but yet above the equipment noise.

We show that changes in access resistance due to incompletely ruptured membrane or other factors, such as mechanical vibration causing the membrane to move with respect to the pipette, can be well measured and compensated (Fig. 6). Hence our approach can be implemented to estimate the excitatory and inhibitory inputs of a cell in these realistic conditions.

Another aspect that might reduce the sensitivity of our method is the presence of pipette stray capacitance. We developed a modular component in the analysis that can be used to correct some of this stray capacitance (Fig. 7). Importantly, no additional measurement is needed beyond the injected sine waves, done in a single trial, in order to measure this stray capacitance and compensate for its effect. Yet, when stray capacitance is much higher than was demonstrated here, this approach fails in providing good estimation of the synaptic conductance. Hence, special care will still be needed to minimize any stray capacitance as much as possible .

We demonstrate in simulations of morphologically realistic neurons, that we can estimate proximal synaptic inputs in a single trial using our approach. Although when compared to simulated voltage-clamp experiments we underestimated these inputs, their shape and relationships were preserved in our measurements as long as the inputs impinged on dendrites not more distant than 300 µm from the soma of our implementation of a pyramidal cell. Hence, this limitation should be considered in real recordings, but also suggest that the method will be able to provide an adequate assessment of proximal inputs .

### Feasibility of the technique in real recordings

Clearly, this framework has to be tested in real recordings of neurons. We fully disclose that we made attempts to test the method in real recordings and discovered that in most of our recordings, none shown here, measurements were unsuccessful. Following tests for impulse response of the amplifier, we found that this results from an active feedback circuit in our intracellular amplifiers. We are currently improving the amplifier circuitry and in parallel developing algorithms that will incorporate the frequency response characteristics of these amplifiers.

### Possible application of the method for measurement of non-synaptic intrinsic conductances

Our method can be used also when voltage-dependent conductances are evolved naturally, as we can measure these inputs at the resting potential of the cell, as long as the sinusoidal fluctuations across the membrane due to the injected current are small. Such an approach therefore can be used when performing pharmacological tests, such as testing effects of modulators, agonists and antagonists of various ion channels. Due to the ability to measure these inputs in a single trial, the time course of the effects can be studied in rapid time scales while examining the effects of such drugs on both inputs at the same time.

In summary, our theoretical study shows that synaptic and other conductances can be measured at high temporal resolution in a single trial when cells are recorded at their resting potential. More research is needed to find if this approach can be used successfully during physiological recordings from real neurons.

## Methods

### Simulations

To develop the method we constructed a simple simulation of a single compartment neuron attached to a resistor, simulating the resistance of the recording pipette. (*R*_*s*_ is the electrode resistance). *I*_*m*_ is the injected current and the other variables as shown in equation 1 and 2. Also note that the capacitive current is given by: *I*_*c*_ = *I*_*m*_ − *k* (*V*_*p*_ − *V*_*m*_)/*R*_*s*_, where *V*_*p*_ is the recorded voltage (across the recording the pipette), *V*_*m*_ is the voltage across the membrane only *I*_*c*_ is stray current. For k = 0 we assume no stray capacitance and for k =1, capacitance was included. Hence:

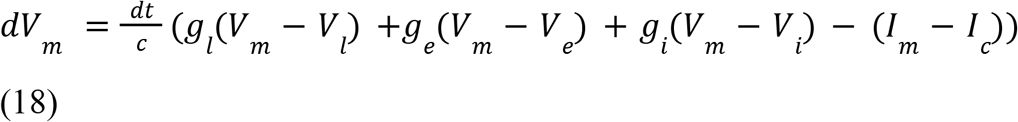

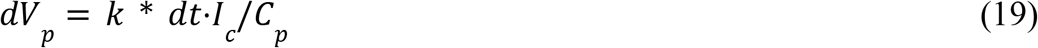

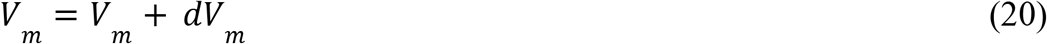

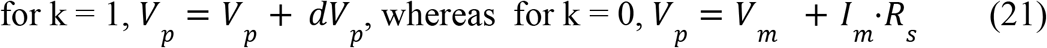

To test the performance of our method in extraction of excitatory and inhibitory conductances, we simulated the response of a cell to a train of synaptic inputs which depress according to the mathematical description of short term synaptic depression (STD,^32^) with *τ*_*inact*_ = 0.003*S* (inactivation time constant) for excitation and *τ*_*rec*_ = 0. 5*S* (recovery time constant) for excitation and the same inactivation time constant for inhibition (0.003S) but a longer recovery time constant (*τ*_*rec*_ = 1. 3*S*) buth exhibiting the same utilization (0.7). The values of the passive properties of the cell and the strengths of synaptic conductances in the simulation were chosen to be at a similar range of experimental data ^8,14,15^. Namely, resting input resistance of 150MΩ, total capacitance of 0.15nF and pipette resistance of 30MΩ. Simulations were run using a simple Euler method with a time step of 0.1 millisecond for all point neuron simulations except for figure 7 (0.025ms).

### Morphologically Realistic Simulations

We used NEURON 7.6.7^33^ in Python 3.7.6 to simulate a CA1 pyramidal cell ^31^. We loaded this cell directly into NEURON without changes to the neuron model. 50 inhibitory and 50 excitatory were distributed on parts of the apical tree. The synaptic mechanism was a modified version of the Tsodyks-Markram synapse ^32^ where we added a synaptic rise time (NEURON mechanism available at https://github.com/danielmk/ENCoI/tree/main/Python/mechs/tmgexp2syn.mod). The synaptic parameters are detailed in Table 1. Event frequency of both synapses was 10Hz and events were jitter with a gaussian distribution of 10 ms STD.

**Table 1.**
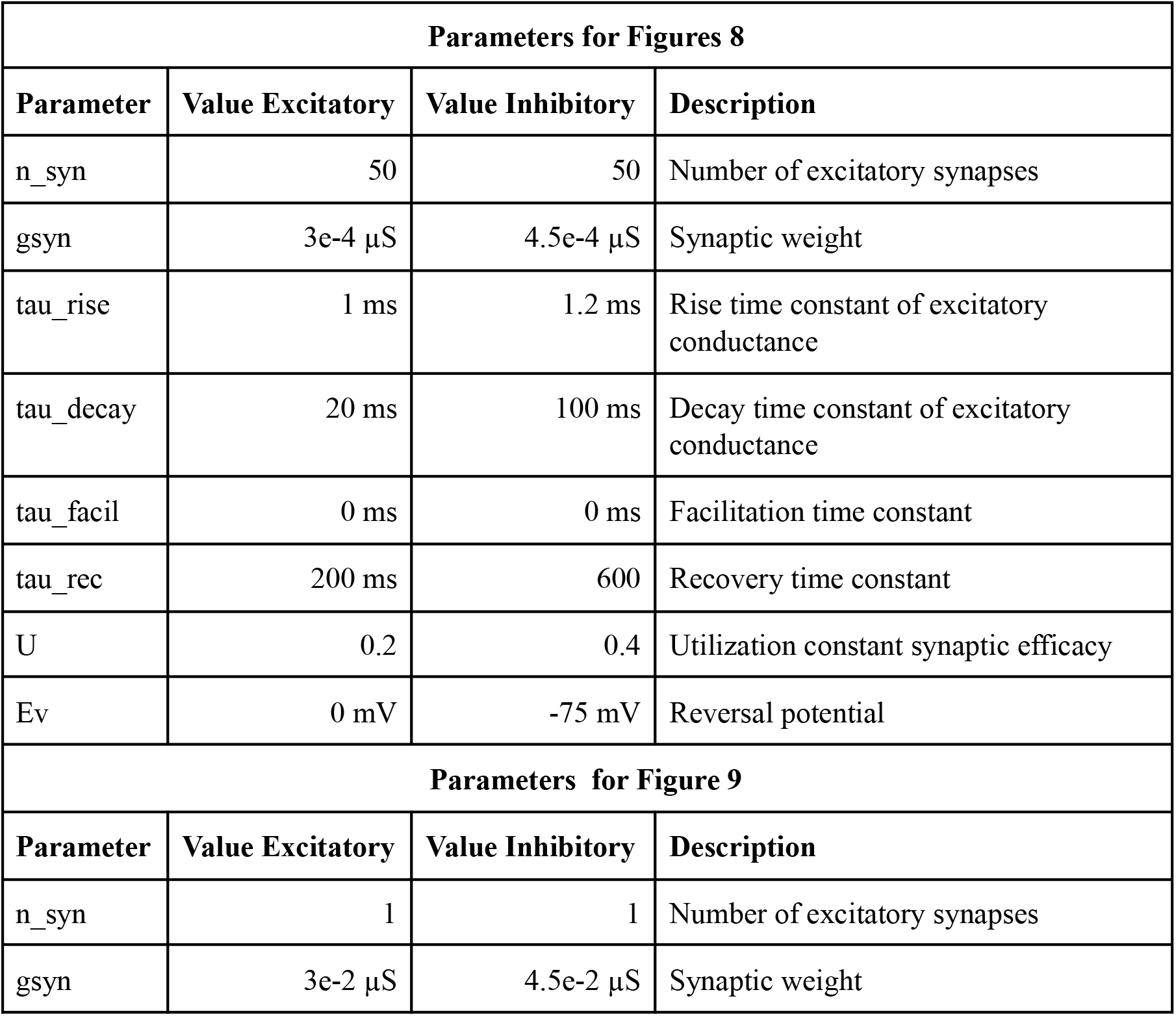
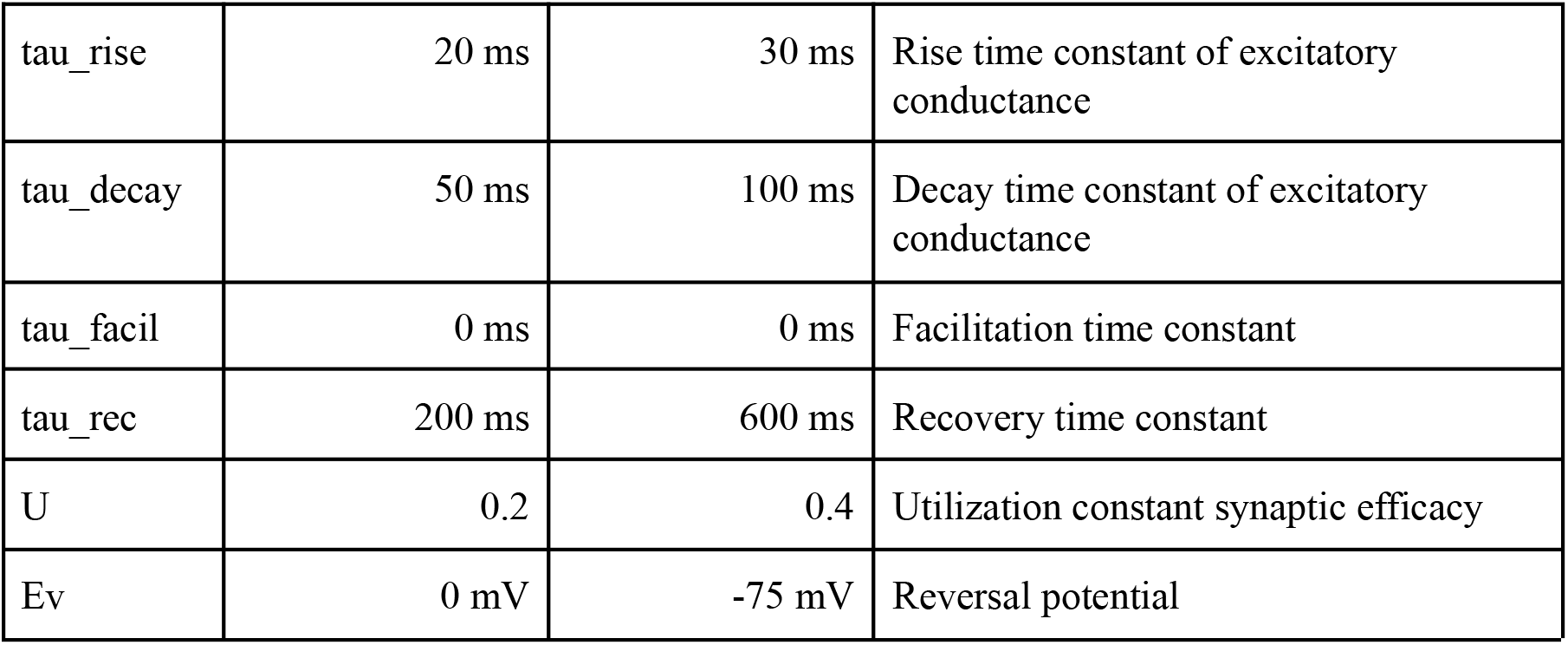
Synaptic parameters of morphologically realistic simulations in Figures 8 & 9.

| Parameters for Figures 8 |  |  |  |
| --- | --- | --- | --- |
| Parameter | Value Excitatory | Value Inhibitory | Description |
| n_syn | 50 | 50 | Number of excitatory synapses |
| gsyn | 3e-4 μS | 4.5e-4 μS | Synaptic weight |
| tau_rise | 1 ms | 1.2 ms | Rise time constant of excitatory conductance |
| tau_decay | 20 ms | 100 ms | Decay time constant of excitatory conductance |
| tau_facil | 0 ms | 0 ms | Facilitation time constant |
| tau_rec | 200 ms | 600 | Recovery time constant |
| U | 0.2 | 0.4 | Utilization constant synaptic efficacy |
| Ev | 0 mV | -75 mV | Reversal potential |
| Parameters for Figure 9 |  |  |  |
| Parameter | Value Excitatory | Value Inhibitory | Description |
| n_syn | 1 | 1 | Number of excitatory synapses |
| gsyn | 3e-2 μS | 4.5e-2 μS | Synaptic weight |
| tau_rise | 20 ms | 30 ms | Rise time constant of excitatory conductance |
| tau_decay | 50 ms | 100 ms | Decay time constant of excitatory conductance |
| tau_facil | 0 ms | 0 ms | Facilitation time constant |
| tau_rec | 200 ms | 600 ms | Recovery time constant |
| U | 0.2 | 0.4 | Utilization constant synaptic efficacy |
| Ev | 0 mV | -75 mV | Reversal potential |

All measurements were performed at the soma. To simulate an access resistor in current clamp we added a section with a specified resistance between the current clamp point process and the soma. The access resistance was 10 MOhm. For the stimulation current we summed two sine waves of 410 Hz and 587 Hz. The combined sine waves had a peak to peak amplitude of 1 nA. Voltage clamp was performed in separate simulations with 10 MOhm access resistance as during current clamp. While isolating the excitatory current, we clamped at the reversal potential of inhibitory synapses (−75 mV). While isolating the inhibitory current, we clamped at the reversal potential of excitatory synapses (0 mV). To convert current to conductance, we divided the current by the clamped voltage minus the synaptic reversal potential.

To investigate the relationship between measurement quality and dendritic path distance to soma, we moved a single excitatory and a single inhibitory synapse to the same dendritic section. Sections were chosen by iterating through the list of apical dendrites in steps of 5. The synaptic parameters are detailed in Table 1.

Python simulation results were saved as .m files using SciPy (^34^. Simultaneous conductance analysis and plotting were performed in MATLAB.

## Data Availability and Code Sharing

All code and most of the simulation results are available at https://github.com/danielmk/ENCoI. The data not included in the repository can be simulated with the provided code.

## Acknowledgements

I would like to thank Hagay Famini and Gal Elyasaf for checking and testing some computational aspects that were used in this study and to Michael Sokoletsly for outstanding comments on the manuscript. I thank Dr. Michael Okun for his comments on the early version of the manuscript and for providing the simulated data used in Figure 6. This work was supported by grants from the DFG-SFB 1089, 01EW1606 -DeCipher EraNet Neuron, HFSP, Israel Science Foundation (ISF 1539/17 and ISF Bikura 2799/20) and Minerva. I.L. is the incumbent of the Norman and Helen Asher Professorial Chair.

## Author Contributions

I.L. developed the method and performed the point neuron simulations. D.M.K. improved the theory and performed the realistic simulation of pyramidal cells. A.P., Y.K. and G.E. provided critical comments during the development of the method and together with D.M.K. tested the feasibility of the method in realistic recordings. H.B. provided crucial comments on the project. I.L. and D.M.K. drafted the paper and wrote it. All authors commented on the results and text.

